# Pupil dilation predicts individual success in emotion regulation and dietary self-control

**DOI:** 10.1101/2020.11.19.376202

**Authors:** Silvia Maier, Marcus Grueschow

**Affiliations:** Zurich Center for Neuroeconomics, Department of Economics, University of Zurich; Neuroscience Center Zurich, University of Zurich, Swiss Federal Institute of Technology Zurich; Translational Neuromodeling Unit, Institute for Biomedical Engineering, University of Zurich and ETH Zurich

**Author notes:** Correspondence should be addressed to, University of Zurich, Department of Economics, Bluemlisalpstrasse 10, 8006 Zurich, Switzerland. denotes equal contribution.

**Keywords:** Pupillometry, arousal, emotion regulation, dietary health, self-control, mood disorder

## Abstract

Multiple theories have proposed that increasing central arousal through the brain’s locus coeruleus – norepinephrine system may facilitate cognitive control and memory. However, for emotion research this hypothesis poses a puzzle, because conventionally, successful emotion regulation is associated with a decrease in arousal.

Pupil diameter is a proxy to infer upon the central arousal state. We employed an emotion regulation paradigm with a combination of design features that allowed us to dissociate regulation- from stimulus-associated arousal in the pupil diameter time course of healthy adults. A pupil diameter increase during regulation predicted individual differences in emotion regulation success beyond task difficulty. Moreover, the extent of this individual arousal boost predicted performance in another self-control task, dietary health challenges. Participants who harnessed more regulation-associated arousal during emotion regulation were also more successful in choosing healthier foods. These results suggest that a common arousal-based facilitation mechanism may support an individual’s self-control across domains.

## Introduction

Self-control skills enable individuals to align their behaviour with own long-term goals and values. Being able to experience different emotional states and if necessary, adaptively regulate emotional reactions, are core aspects of daily human lives (Gross, 2015; Gross & Barrett, 2013). Disturbances of emotion regulation are a hallmark of multiple psychiatric disorders (Gross & Jazaieri, 2014; Kring & Sloan, 2010; Zilverstand, Parvaz, & Goldstein, 2017) and emotion control has recently been identified as the component of self-control that is most predictive of mental health (Eisenberg et al., 2019). For example, individuals who engage more automatically in reappraisal when encountering negative experiences may mitigate stress-inducing effects (Thoern, Grueschow, Ehlert, Ruff, & Kleim, 2016) through this emotional buffer (Shahane, Lopez, & Denny, 2018). Despite the abundance of psychological disorders involving maladaptive emotion regulation (World Health Organization, 2017), validated quantitative tools to assess an individuals’ engagement in regulation in a clinical or research setting are surprisingly scarce and urgently needed (Kalisch et al., 2017).

Novel frameworks conceptualize how emotion regulation operates on a moment-to-moment basis (Kalisch, 2009; Sheppes & Gross, 2011) and across biological and psychological levels. However, it is not trivial to measure whether individuals engage in regulating their emotions at any given moment, and most importantly, to predict how successful they will be. These questions are paramount for both basic and applied research, because inflexibility or inability to adaptively regulate emotions through strategies that favour beneficial behaviour in the long term is a hallmark of diseases such as depression, eating disorders, substance abuse, and posttraumatic stress disorder (Joormann, Yoon, & Siemer, 2010).

We therefore aimed to simultaneously quantify the onset and efficacy of human emotion regulation. To this end, we combined an established emotion regulation paradigm (Buhle et al., 2014; Denny, Ochsner, Weber, & Wager, 2014; Goldin, McRae, Ramel, & Gross, 2008; Ochsner, Bunge, Gross, & Gabrieli, 2002; Ochsner et al., 2004; Ochsner, Silvers, & Buhle, 2012; van Reekum et al., 2007; Wager, Davidson, Hughes, Lindquist, & Ochsner, 2008) with pupillometry – a measure intricately linked to activity in the arousal system (Joshi, Li, Kalwani, & Gold, 2016; Varazzani, San-Galli, Gilardeau, & Bouret, 2015; Zerbi et al., 2019). Its millisecond temporal resolution allowed us to assess at which time point individuals engaged in emotion regulation. One crucial problem for the interpretation of such physiological readouts in emotion regulation research is whether an increase in pupil dilation relates to stimulus-associated arousal or to engagement in genuine cognitive control processes. To solve this question, we employed a combination of design features in the emotion regulation paradigm that allowed us to separate regulation- from stimulus-associated arousal components in the pupil diameter time course. Moreover, we aimed to predict from individual pupil diameter during the regulation period to which degree participants managed to render their emotions more neutral. Our findings present a physiological account based on pupil dilation that quantifies an individual’s engagement and success in emotion regulation. Furthermore, we additionally test whether this measure captures features of an individual’s regulation ability that generalize to another self-control domain (Duckworth & Tsukayama, 2015), namely dietary health challenges.

That the pupil dilates in response to emotionally relevant stimuli has been corroborated by ample evidence (Bradley, Miccoli, Escrig, & Lang, 2008; Bradley, Sapigao, & Lang, 2017; Ferrari et al., 2016; Henderson, Bradley, & Lang, 2014, 2018; Hess & Polt, 1960; Kret, Roelofs, Stekelenburg, & de Gelder, 2013; Kuntz, 1929; Partala & Surakka, 2003; Snowden et al., 2016). Individual differences in pupil dilation in response to emotional stimuli show excellent test-retest reliability (Hess & Polt, 1960; National Advisory Mental Health Council Workgroup on Tasks and Measures for Research Domain Criteria, 2016). Pupillometry allows quantifying the pupil dilation response precisely, and pupil dilation and constriction have been related to activity in the sympathetic and parasympathetic branches of the autonomic nervous system (Loewenfeld, 1958; Lowenstein & Loewenfeld, 1950a, 1950b, 1958). Lesion studies show that the parasympathetic pathway primarily controls the pupil light reflex, i.e. pupil constriction in response to brightness changes, whereas the sympathetic pathway controls dilation in response to arousal (Loewenfeld, 1958). Arousal could for example be caused by emotionally salient stimuli, but also be recruited in order to meet cognitive demands (Aston-Jones & Cohen, 2005; Eldar, Cohen, & Niv, 2013, 2016; Mather, Clewett, Sakaki, & Harley, 2016; Verguts & Notebaert, 2009) or in order to mobilize effort (Kurniawan, Grueschow, & Ruff, 2020; Varazzani et al., 2015; Walton & Bouret, 2019). Previous studies have also reported that pupil diameter increases when individuals regulate their emotions (Johnstone, van Reekum, Urry, Kalin, & Davidson, 2007; Kinner et al., 2017; Richey et al., 2015; Urry, 2009; Urry et al., 2006; van Reekum et al., 2007).

During mental problem solving, pupil dilation is strongly correlated with the difficulty of the problem, as Hess and Polt (1964) first described. They suggested a measure of “total mental activity”, reflected in a combination of the latency and the amplitude of the pupil response, that we mimic in our approach. Kahneman and Beatty (1966) elegantly demonstrated how the pupil dilates in response to processing load and constricts once the load is reduced (see Beatty (1982) for review of earlier works and van der Wel and van Steenbergen (2018) for a recent review on pupil dilation during cognitive control).

Therefore, we assume that the pupil dilation signature during emotion regulation is informative about the individual cognitive engagement during this self-control process. We chose to investigate the engagement in regulation via pupil dilation because the ability to engage self-control is fundamental for how well individuals can follow through on their goals, which is hard to assess in an unbiased fashion via self-report. Because the participant cannot covertly manipulate pupil dilation, it provides an unbiased readout of the cognitive engagement in self-control, whereas self-reports have been shown to suffer from several subjective biases (DeVylder & Hilimire, 2015; Logan, Claar, & Scharff, 2008). For elementary cognitive control processes such as updating, attention shifting, action inhibition and explore-exploit trade-offs, an adaptive modulation via arousal has been reported and measured using pupil dilation (see van der Wel and van Steenbergen (2018) for review). We thus hypothesized that individuals who increase their cognitive engagement and hence arousal levels to best meet the task requirements should be more successful regulators. We set out to test whether individuals who are able to boost task-relevant, adaptive processing through arousal (quantified here using pupil dilation) achieve higher levels of emotion control. In our experiment, we demonstrate that this relationship holds for emotion regulation, and critically also transfers to another complex behavioural task, namely solving dietary health challenges. These two tasks employ important mechanisms associated with a wide array of disorders of affect and interpersonal conduct, obesity, and addiction (Fernandez, Jazaieri, & Gross, 2016; Lempert, Steinglass, Pinto, Kable, & Simpson, 2019).

Using the pupil dilation signal, we predict how successful individuals apply an instructed regulation strategy, reappraisal. To provide a generalizable account, we investigate commonalities in the process of regulating emotional experiences of positive and negative valences. We show that the pupil dilation index predicts individual differences in regulation success. Moreover, we demonstrate its convergent validity, as the pupil dilation index measured in the emotion regulation task is associated with individual regulation success in a separate dietary health challenge. This strongly suggests that the pupil dilation index captures a more general process supporting adaptive behaviour and regulation success across multiple task domains and may serve as a promising measure in future investigations of self-control mechanisms in health and disease.

## Results

### Emotion reappraisal task

The fMRI part of this experiment was used to test a separate hypothesis that is reported in Maier and Hare (2020). For completeness and easier accessibility of the current paper, we summarize the behavioural results on emotional stimulus reappraisal again briefly in the Supplementary Information. After viewing or reappraising affective pictures, participants rated their current affective state. When we modelled emotion ratings as a function of block type, the results showed that participants successfully reappraised the emotional content of the images (Supplementary Methods, Supplementary Results, Supplementary Table 1, and Figure 1a reproduced from data published in Maier and Hare (2020)).

**Figure 1.**
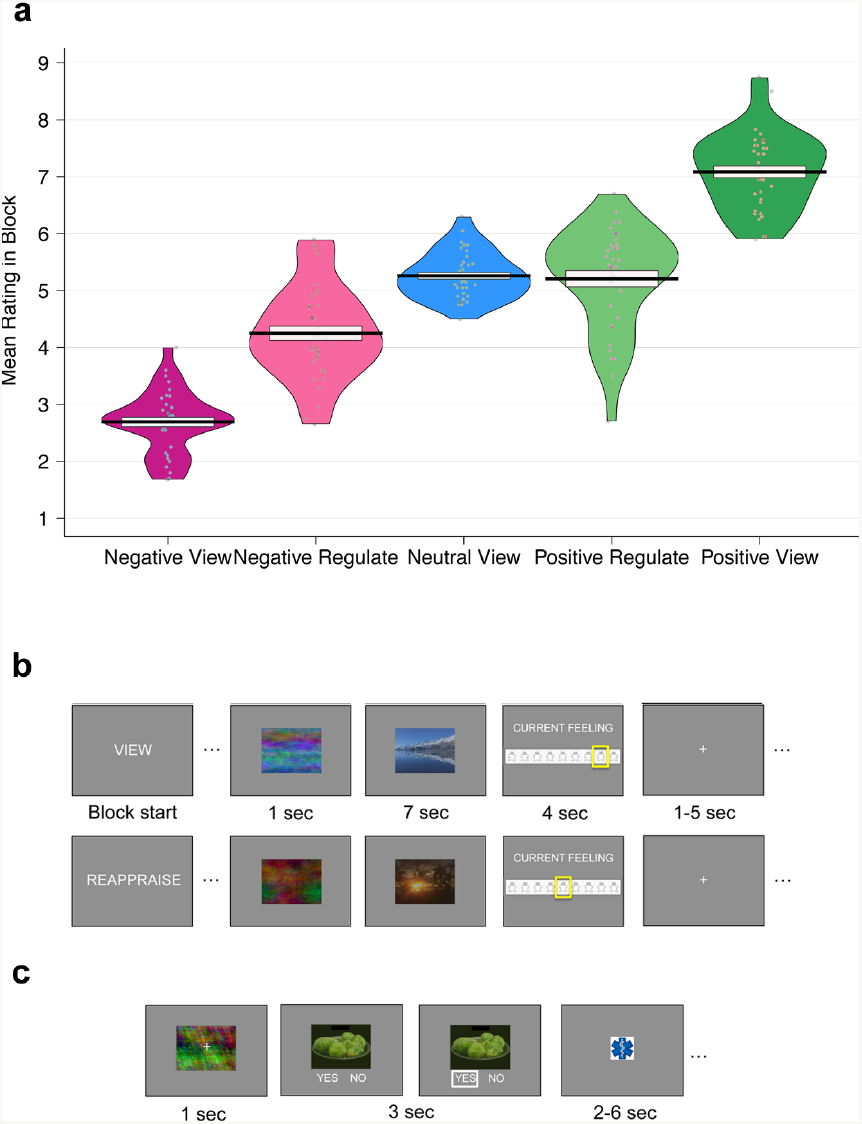
Self-regulation tasks and behaviour. **a)** Emotion reappraisal success. On the 9-point SAM scale, the panel shows the mean emotion ratings for each block (negative view, negative reappraise, neutral view, positive reappraise, and positive view blocks). The black solid line indicates the group mean and the box its standard error. Each grey dot represents the mean ratings from one participant. Participants successfully reappraised both negative and positive images, shifting their feelings in both cases towards a neutral state. **b)** Emotion reappraisal task. Participants saw positive, neutral and negative stimuli from the International Affective Picture System (IAPS). For display purposes, we replaced the IAPS stimuli by unrelated landscape photos here. In each block, participants either viewed the images without changing their emotional response (“view”) or reappraised the scene to render their feelings more neutral (“reappraise”). To allow the pupil to adapt to brightness and contrast, a phase-scrambled version of the stimulus was displayed for 1 second before the image was revealed. Participants then viewed or reappraised the scene for 7 seconds before they rated their current feeling on a Self-Assessment-Manikin (SAM) scale (for details see Methods). A jittered inter trial interval of 1-5 seconds separated the trials. **c)** Dietary health challenge task. Participants made 100 choices indicating whether they wanted to eat the displayed food at the end of the study. A phase-scrambled adaptation stimulus was presented for 1 second before the food image was revealed and participants had 3 seconds to decide by pressing the left or right button for selecting the answers “yes” or “no”. A white frame highlighted the answer for 0.1 seconds when it was logged. Trials were separated by a 2-6 second (jittered) inter-trial interval.

### Pupil during emotion regulation

To test our hypothesis that regulation signals are reflected in the pupil dilation signature, we compared the baseline-corrected pupil dilation time course during reappraisal with the pupil dilation time course in the view condition. Positive and negative reappraisal showed similar pupil dilation differences over time: Regulation was characterized by an increase in pupil diameter from around 2 seconds onwards, compared to the viewing conditions, in which the pupil started constricting on average again at this time (Figure 2a). Hence the additional pupil dilation during reappraisal may indicate participants are engaging in the regulation process, regardless of valence.

**Figure 2.**
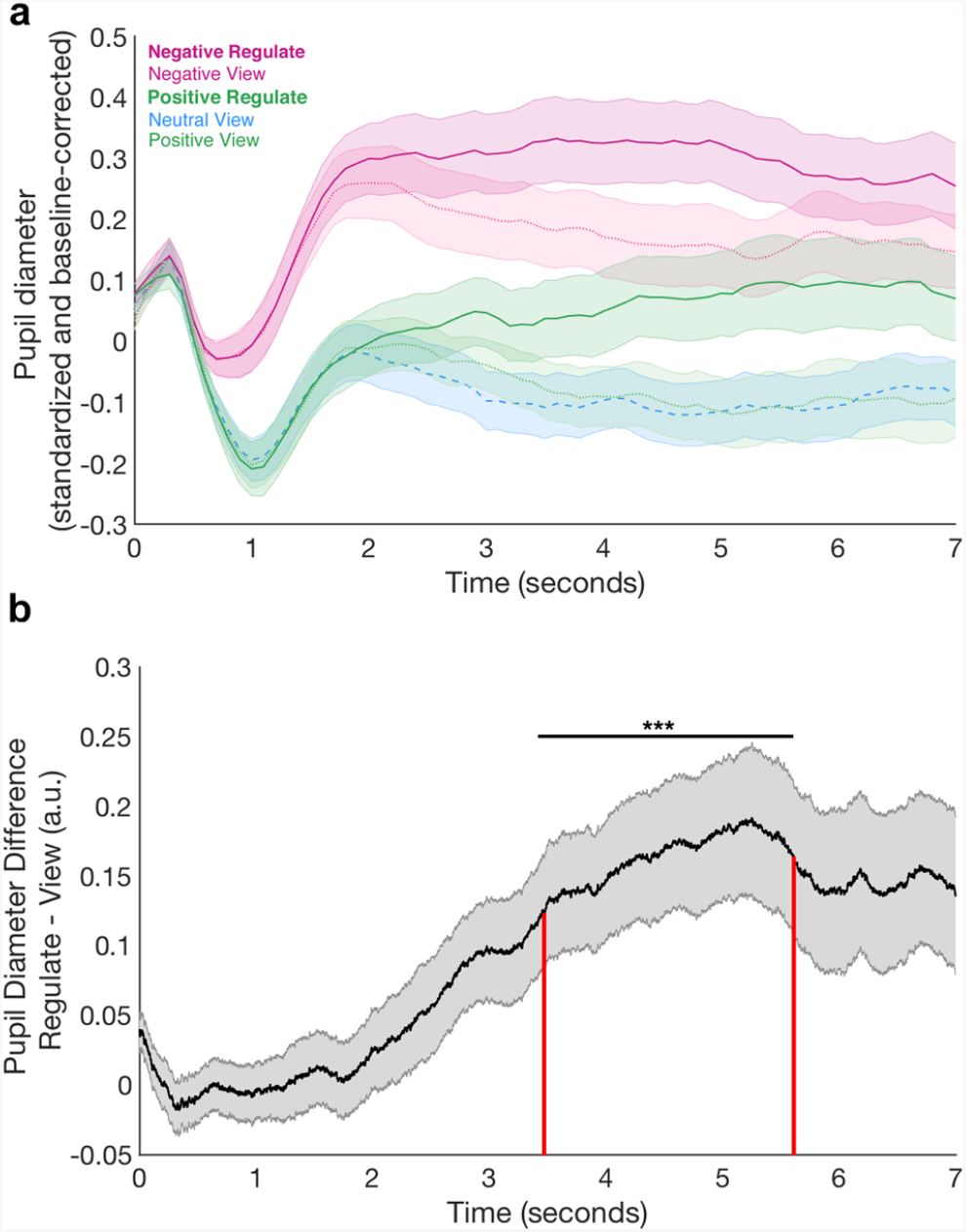
Pupil results. **a)** Mean pupil dilation during the 7-second regulate (saturated colours) and view periods (lighter colours). The signal across the whole emotion task was z-scored within-participant and baseline-corrected on each trial for the pupil size in the 500 milliseconds prior to the regulation period (during which the phase-scrambled adaptation version of the stimulus was displayed, see Methods). The mean pupil dilation was calculated over all 20 trials in each block for each participant and then averaged across the group. The shaded areas indicate the standard error of the mean across participants. Lines denote the mean pupil dilation at each time point. The dotted blue line represents the neutral view condition for comparison. **b)** Pupil Diameter Difference between Regulation and View Trials. We collapsed over positive and negative blocks in order to test for a valence-independent regulation signal across all participants. The mean z-scored pupil dilation for positive and negative view trials was subtracted from the mean z-scored dilation during positive and negative reappraise trials for each participant. A cluster-based permutation test indicated that between 3.4 and 5.6 seconds of the reappraisal/view period, the pupil dilation in reappraisal blocks was larger than during viewing blocks (p < 0.001; indicated by the black horizontal line and stars). For each participant, the mean of the Reappraisal – View difference in the pupil dilation signal during this period (marked by the red vertical lines in the plot) determines the pupil dilation index. The shaded areas indicate the standard error of the mean across participants. The black graph denotes the mean pupil dilation at each time point.

In order to rigorously test during which time period pupil dilation signatures for regulation and viewing differed without having to predefine a window for the analysis, we used a cluster-based permutation t-test. This method identified adjacent time bins in the pupil time course that significantly differed between the regulation and view conditions. To characterize a valence-independent regulation signature, we collapsed the data across positive and negative regulation periods and generated a contrast that identified changes due to regulation regardless of valence. For each participant, we averaged the pupil time course during positive and negative reappraisal and subtracted the average pupil time course during positive and negative view conditions. We then performed a non-parametric cluster-based permutation t-test on this contrast. This analysis confirmed that the pupil was dilated more during regulation compared to the viewing periods: Within the stimulus presentation time window of 7 seconds, the test indicated that between 3.4 and 5.6 seconds after stimulus onset the pupil was significantly more dilated during regulation than during viewing (Figure 2b; mean PDI across the group = 0.16 ± 0.003, p < 0.001, maximum T-value during this time = 3.74). We thus identified the time period during which we could reliably measure a regulation signal in the pupil time course.

As we had subtracted out all activity related to viewing pictures of equivalent emotional valence and arousal, and had controlled the physical stimulus properties in the analysis, we reasoned that during this time window, the pupil dilation difference between the reappraise and view condition would most likely be driven by cognitive processes supporting regulation (or concomitant increases in arousal). Notably, we can exclude the possibility that the signal was merely driven by the natural arousal level usually created by these stimuli, because we had designed the blocks such that stimuli shown in the regulation and view conditions were equated for their average arousal ratings, and thus the average arousal pertaining to the stimuli was individually controlled for when generating the contrast. We can also exclude any other properties of the stimulus sets driving this effect, because the stimulus sets in each condition featured equally often on both sides of the subtraction contrast. We thus calculated the mean difference of the Reappraise > View contrast during the significant time window in order to quantify individual differences in regulation engagement and used this measure as Pupil Dilation Index (PDI) of regulation.

### Pupil predicts regulation success

Using this pupil dilation index, we performed a Bayesian linear regression (Eq. 3a) to account for variations in emotion regulation success between individuals. As task difficulty may impact on regulation success, the regression included the average difficulty each participant faced. As a proxy for difficulty, we calculated a measure for the affective distance of the stimulus from neutral. The absolute distance between the post-task view ratings for the regulated images and the neutral point of the rating scale constitutes the affective distance. The mean of the affective distance over all trials generates one index of average task difficulty for each participant. Both the pupil dilation index (Figure 3a, Table 1; beta = 0.34 ± 0.14 Standard Deviation (SD), 95% Credible Interval (CI) = [0.06; 0.61]) and affective distance (beta = 0.30 ± 0.14 SD, 95% CI = [0.02; 0.58]) explained substantial portions of variance in regulation success across participants. Greater central arousal activity during regulation was thus related to greater reappraisal success, above and beyond the effects of affective distance.

**Table 1.** Emotion regulation success explained by pupil dilation index and affective distance.

| a) | Beta Estimate | Standard Deviation | 95% Credible Interval |
| --- | --- | --- | --- |
| (Intercept) | 1.77 | 0.14 | [1.50; 2.03] |
| Pupil Dilation Index | 0.34 | 0.14 | [0.06; 0.61] |
| Affective Distance | 0.30 | 0.14 | [0.02; 0.58] |
| Pupil Dilation Index X Affective Distance | 0.21 | 0.19 | [-0.15; 0.58] |
| b) |  |  |  |
| (Intercept) | 1.83 | 0.20 | [1.43; 2.24] |
| Pupil Dilation Index | 0.33 | 0.16 | [0.01; 0.64] |
| Affective Distance | 0.28 | 0.15 | [-0.01; 0.57] |
| Age | 0.07 | 0.15 | [-0.23; 0.35] |
| Task order | -0.13 | 0.29 | [-0.71; 0.44] |
| Pupil Dilation Index X Affective Distance | 0.20 | 0.20 | [-0.18; 0.61] |

**a)** Results from the Bayesian Linear Regression specified in Eq. 3a. Regulation success (measured on the 9-point SAM scale) was modelled by the mean-centred and standardized coefficients for Pupil Dilation Index (measured as mean difference in the pupil dilation curve for the Regulate > View contrast in the significant regulation time between 3.4 and 5.6 seconds) and Affective Distance (measured as the absolute value of the distance of the view rating from the neutral valence on the SAM scale). Pupil Dilation Index and Affective Distance were interacted, to test whether it might be easier or harder to reappraise with smaller or greater affective distances.
**b)** Augmenting the model (Eq. 3b) by the control regressors for Age and Task Order shows that the explanation of regulation success by PDI is robust to controlling for the age and the order in which participants completed the emotion regulation and dietary health challenge tasks. Model fits are given as the population level mean of the posterior distribution ± standard deviation (SD) and the 95% Credible Interval.

**Figure 3.**
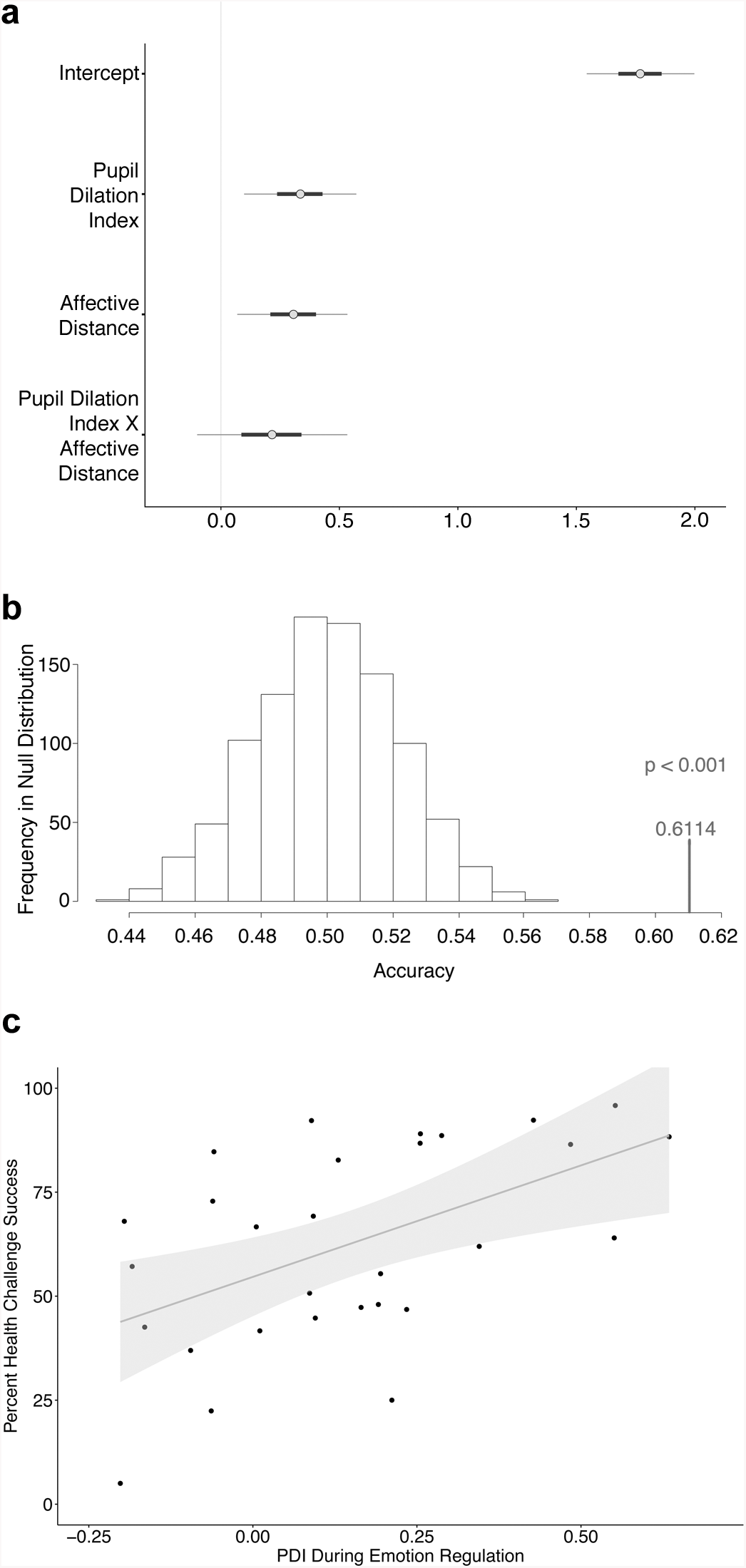
Predictive and convergent validity of the pupil dilation index for self-regulation success. **a)** Bayesian linear regression of self-regulation success using the model given in Eq. 3a. Pupil Dilation Index denotes the coefficient estimating the influence of regulation-related arousal components. Affective Distance denotes the coefficient estimating the influence of the mean absolute difference of the view rating from neutral (middle of the rating scale) that can be interpreted as a proxy for regulation difficulty. Regulation success increased both with greater Pupil Dilation Index and Affective Distance. The plot shows the mean beta estimates (grey dots) as well as the range of coefficients within the 90% Credible Interval) that is shown by the light grey horizontal bars (thick horizontal bars = 50% Credible Interval). **b)** Predicting regulation success out-of-sample. The results from the model in Eq. 3a were cross-validated with a Leave-2-Participants-Out approach. Based on data from N-2 participants, we predicted which of the two left-out individuals regulated more successfully. The model predicted with 61% accuracy significantly above chance (p < 0.001). **c)** Convergent validity of the Pupil Dilation Index. The pupil dilation index (PDI) that we measured during emotion regulation also shows convergent validity across different types of self-control tasks: A higher PDI value was associated with higher health challenge success scores within-individual in the dietary health challenge task that individuals completed in the same experimental session (r = 0.51, posterior probability rho > 0 = 0.9995), explaining 26% of the variance (R-squared = 0.26) in dietary health challenge success. The grey shaded area signifies the 95% confidence interval.

In order to check the robustness of this result, we ran a control analysis augmenting the model by the age of the participants and the order in which they performed the emotion regulation and dietary health challenge tasks (Eq. 3b). When accounting for these alternative explanations of regulation success scores, the pupil dilation index still explained signification portions of the variation in reappraisal success (PDI coef. = 0.33 ± 0.16 SD, 95% CI = [0.01; 0.64]), whereas none of the control regressors did (Table 1b). In order to rule out that the result was driven by potential reappraisal spillover effects on the view ratings that were taken after the images had been first reappraised, we also ran a control analysis for the model in Eq. 3a using block-wise measures for reappraisal success and affective distance that we constructed based on the equivalent stimuli from the “view” condition and found that our conclusions remained unchanged (see Supplementary Methods, Results and Discussion; Supplementary Figure 2 and Supplementary Table 2).

To test the predictive validity of the pupil dilation index, we first conducted a leave-two-participants-out analysis, in which we fit the model specified in Eq. 3a to a training set of all combinations of participants leaving out two participants as a test set. On each iteration we predicted based on the estimates of the training set for the two test participants which of two individuals would regulate more successfully. We successfully predicted with 61.14% accuracy out-of-sample which of the test participants was better at regulating emotion responses (Figure 3b). A permutation test revealed that this accuracy would have occurred on less than 1 in 1000 occasions if the prediction were based on randomly labelled instead of true data (p < 0.001).

### Convergent validity across self-control domains

In order to test the convergent validity of the pupil dilation index across self-control domains, we performed a Bayesian rank correlation analysis between the pupil dilation index from the emotion regulation task and the health challenge success levels that the same individuals achieved in a separate dietary health challenge task completed on the same day. A higher pupil dilation index during emotion regulation was associated with higher dietary health challenge success levels in this separate task (Spearman’s rho = 0.51, 95% Credible Interval = [0.232; 0.759], posterior probability (rho > 0) = 0.9995; Figure 3c; R-squared = 0.26). Critically, this association was robust to the order of the self-regulation tasks (Eq. 4). Task order did not explain variation in dietary health challenge success levels beyond the pupil dilation index (Table 2).

**Table 2.** Dietary health challenge success by pupil dilation index and task order.

| Population-level effects | Beta Estimate | Standard Deviation | 95% Credible Interval |
| --- | --- | --- | --- |
| (Intercept) | 61.96 | 5.81 | [50.88; 73.89] |
| Pupil Dilation Index | 12.47 | 4.40 | [3.39; 21.08] |
| Task Order | 1.39 | 8.48 | [-15.58; 17.81] |

*Dietary health challenge success modelled with a Bayesian linear regression as a function of pupil dilation index and task order (Eq. 4): The individual dietary health challenge success level (measured over all challenging trials in percentage points) is explained by the standardized and mean-centred Pupil Dilation Index (PDI). Compared to the average level of PDI in the group, an increase of 1 standard deviation (SD) in pupil dilation explains an additional 12.47% of dietary health challenge success on top of the 61.96% health challenge success that an individual with average PDI would show. The analysis is controlling for a factor representing Task Order (emotion regulation or dietary choice task first). Task order does not explain variation in dietary health challenge success levels (95% Credible Interval for the estimate includes zero).*
*Model fits are given as the population level mean of the posterior distribution ± standard deviation (SD) and the 95% Credible Interval.*

The link of the pupil dilation index to overall success levels in both emotion regulation and dietary health challenges suggests that the pupil dilation index captures a more general process underlying self-regulation success in both task domains. Its predictive power with 26% explained variance across tasks (R-squared = 0.26) provides converging evidence that the pupil dilation index measured an important correlate of self-control success.

## Discussion

We find that the pupil dilation difference between reappraising and viewing emotional stimuli predicts the degree to which individuals are able to render their feelings more neutral. We demonstrate that this is tied to the individual level of regulation-related arousal, indexed by pupil dilation, which we separate from the estimated effects of task difficulty that we quantify via affective distance (Eq. 3a). This finding was independent of the participant’s age and the order in which the self-control tasks were performed (Eq. 3b). In order to ensure that the relationship was not merely driven by potential spillover of reappraisal on the subsequently collected ratings under view conditions (MacNamara, Ochsner, & Hajcak, 2010), we performed a control analysis with measures constructed based on the “view” condition. Note that any reappraisal spillover would have weakened the evidence instead of supporting our conclusions. In the control analysis, we found the same benefit of higher regulation-associated arousal to emotion regulation success (see Supplementary Methods and Results).

It also speaks against an explanation of the regulation success purely by an experimenter demand effect that we identified a relationship between the regulation success and a physiological measure for which voluntary manipulations would have been easily detected. This association should not hold if all variance in the reported emotion after regulation were explained by the desire to conform to the task instructions. As a further piece of evidence that the emotion task and pupil analyses worked as planned, our results for viewing emotional stimuli without regulating replicated previous findings (Bradley et al., 2008; Kinner et al., 2017; Partala & Surakka, 2003; Urry et al., 2006; van Reekum et al., 2007).

Importantly, we isolated regulation-related signals in the pupil dilation time course. The experimental design required to compute the pupil dilation index is rather simple and can be repeated with a broad range of stimuli for other cognitive experiments that go beyond the realm of emotion control. As we demonstrate in our paper, potential interpretative caveats can be readily addressed upfront through experimental design features that allow to construct the pupil dilation index. These are: 1) Collecting norm ratings of the stimulus material in terms of arousal and valence beforehand, based on which 2) the experimental paradigm is constructed such that valence and arousal aspects are equated on average across the condition of interest and control condition, so that 3) any remaining portions of the pupil signature reflect the cognitive process(es) of interest that can be isolated by a subtraction contrast. 4) Adjusting the physical features of the stimuli across the sets for the control condition and the condition of interest in terms of brightness and contrast allows to interpret the subtraction contrast just in terms of the content features, and 5) the 1-second adaptation period with a phase-scrambled stimulus allows to interpret the pupil signal in continuous time from the start of the cognitive process of interest. Based on these prerequisites, 6) using a cluster-based permutation test facilitates tracking the cognitive process of interest in continuous time. Here, we employed a set of emotional stimuli with known valence and arousal norms, which have been successfully used in research through decades, but in principle, a large variety of stimulus material can undergo such norming to mimic our design.

Our results indicate that both the pupil dilation index and affective distance independently explain variance in how successfully individuals render their feelings more neutral, pointing towards separable factors that influence self-control success. Despite subtracting out the signal related to the average arousal properties of the stimuli through the Reappraise > View contrast, we still observed pupil dilation, indicating increased arousal during the process of regulating. If the pupil dilation signature during the regulation condition was purely explained by the arousal caused by viewing the content of the image, the pupil diameter should decrease once regulation sets in. In our regulation condition, however, pupil dilation stayed elevated for a prolonged period, suggesting that regulation still further engaged the pupil-linked arousal system.

The pupil dilation index, a proxy for the central arousal state, was an indicator of how well participants applied cognitive control across tasks. We observed in this study that a higher arousal state seemed to benefit the application of cognitive control, yet we are not proposing a directional relationship at this point: It may either be the case that higher arousal instigates more control, or, vice versa, that during cognitive control, additional arousal is recruited to fine-tune control processes.

One reason for our observations may be that participants engage arousal in order to regulate. This is in line with reports of higher pupil dilation readouts during similar tasks (Urry et al., 2006; van Reekum et al., 2007) that show a similar pattern of increasing sympathetic signalling within the regulation period. Pupil dilation is a natural candidate to measure such arousal in service of cognitive control (Gilzenrat, Nieuwenhuis, Jepma, & Cohen, 2010; Laeng, Orbo, Holmlund, & Miozzo, 2011; Rondeel, van Steenbergen, Holland, & van Knippenberg, 2015; Steinhauer, Siegle, Condray, & Pless, 2004; van der Meer et al., 2010; van Steenbergen & Band, 2013; Vo et al., 2008; Wang, Brien, & Munoz, 2015), because it is tightly coupled to activity in the locus coeruleus that releases noradrenaline (Clewett, Huang, Velasco, Lee, & Mather, 2018; Eldar et al., 2013; Lee et al., 2018; Murphy, O’Connell, O’Sullivan, Robertson, & Balsters, 2014; Verguts & Notebaert, 2008).

Our data are consistent with different mechanistic theories on the relationship between arousal and cognitive control processes. However, our experiment was not designed to distinguish between potential mechanisms (such as effort, attention, or working memory processes) that may all be invigorated by arousal in order to support cognitive control, and all would yield pupil signatures similar to the one we observed.

For example, Varazzani et al. (2015) suggested that noradrenaline release of the LC may serve to increase motivation to exert effort for an encountered challenge at the time when an energetically costly action is performed. Moreover, the “adaptive gain” theory (Aston-Jones & Cohen, 2005) postulated more generally that phasic noradrenaline release by the locus coeruleus helps to facilitate behaviours that optimize task performance. Mather and colleagues have suggested in their GANE theory that arousal may lead to norepinephrine release that facilitates selective attention (Dahl, Mather, Sander, & Werkle-Bergner, 2020) and memory consolidation (Mather et al., 2016). Similarly, Verguts and Notebaert (2009) have proposed in their “adaptation by binding” theory that arousal enhances cognitive control by facilitating communication between task-relevant cortical areas and increasing online Hebbian learning that binds together representations that are activated together. Attentional focus and modification of associations in memory are both relevant to emotion regulation via reappraising the content of the pictures. In line with the “GANE” and “adaptation by binding” theories, one might speculate that the impact of arousal on memory processes may contribute to reappraisal success: If old associations that have been activated were more malleable under arousal, the increase in arousal during the regulation process may foster online learning of new associations (for a review on pupillometry measures of memory formation and retrieval see Papesh and Goldinger (2015)). Moreover, this process may profit from better attentional focus on the aspects of the stimulus that are to be re-framed. Regardless which of these mechanistic explanations contributes to the more successful reappraisal, it is noteworthy that we seem to tap into a more general process that holds beyond the reappraisal of emotional stimuli. The pupil dilation index has predictive validity, as evidenced by the out-of-sample prediction of emotion regulation success. We also demonstrate its convergent validity: it is associated with success in a dietary health challenge task. This strongly suggests that the pupil dilation index captures a more general process supporting successful regulation across multiple task types and may serve as a promising measure in future investigations of self-control mechanisms in health and disease. At this point, we can only speculate which aspect of emotion regulation the pupil dilation index picks up that links to the success in both tasks.

Fernandez et al. (2016) proposed that emotion regulation might form an own domain according to the Research Domain Criteria (RDoC) that emerges when other RDoCs are combined (negative valence systems, positive valence systems, cognitive systems, social processes, and arousal and regulatory systems). Through our experimental approach, we narrow down the common contributors to self-control success across domains that we indexed with the pupil measure to the RDoC components of cognitive systems, and arousal and regulatory systems. We demonstrate a link to self-regulation success across domains specifically through an autonomic nervous system measure used as a proxy for central arousal levels (pupil dilation). We also found in prior work that another index of the flexibility of the reaction of the autonomic nervous system, heart rate variability, predicted health challenge success in dietary choices (Maier & Hare, 2017). We may therefore cautiously speculate that the reactivity of arousal systems might play a role in self-control success through a facilitation mechanism in which dynamic changes in arousal support or tune cognitive processes for self-control. Our results also suggest that a greater dynamic range for increasing central arousal may help to boost emotion control.

Taken together, our study presents several important advances. We introduce an approach to measure regulation effects in pupil dilation in a continuous fashion. This advances the field, because an analysis in pre-defined, coarse-grained time-windows does not allow capturing the actual onset of regulation. With our approach, pupil diameter indices do not have to be aggregated any more over a priori defined time-bins in order to quantify regulation (Bebko, Franconeri, Ochsner, & Chiao, 2011; Kinner et al., 2017; Urry et al., 2006; van Reekum et al., 2007), but can be tracked within the continuous pupil dilation signal. The studies by Kinner et al. (2017) and Urry et al. (2006) provide interesting leads, showing that different regulation strategies express different forms of pupil dilation time courses. This potentially allows tracking through different pupil signatures which type of emotion regulation strategy is applied, instead of relying on more burdensome EEG setups (Thiruchselvam, Blechert, Sheppes, Rydstrom, & Gross, 2011).

A potential application of the pupil dilation index in affective disorders may be to measure the progress an individual makes in finding and applying more effective and appropriate strategies to regulate their reactions to emotionally salient events or thoughts (Gross, 2015; Kross, 2015). We believe that this may foster further lines of research, for instance on how individuals flexibly utilize different types of emotion regulation strategies (Bonanno & Burton, 2013; Sheppes & Levin, 2013; Suri et al., 2018). In addition, our approach might help to assess individual progress in applying trained strategies to reduce for example cravings for foods or drugs of abuse. In the future, our results may be relevant to diagnosis, treatment and intervention in psychiatric disorders and psychosomatic medicine.

In sum, we have presented a combination of innovative approaches that advance both basic and applied research on emotion regulation, and our understanding of self-control and cognitive control mechanisms more generally. We believe that pupillometry can ideally leverage modern technological advances in data acquisition and analysis. Modern eyetrackers are highly mobile and could be used at bedside or in workplace settings. Data acquisition is comparatively cheap, and adjustment of stimulus material and artefact correction have become easier with modern computational tools, yielding potentially better data quality than other typical readouts of the autonomic nervous system such as heart-rate or skin conductance. Measuring pupil dilation is unobtrusive, and it cannot be manipulated covertly by the participant. Combining these advantages enables assessing spontaneous changes in arousal without asking individuals for their self-report. For clinicians, this facilitates data collection when individuals have difficulties with regard to interoception or introspection, and for basic science, it yields neurophysiological readouts to develop and to test theories.

Taken together, our physiological and behavioural data across different types of self-control domains suggest that a common arousal-based facilitation mechanism contributes to individual differences in human self-control in complex tasks. We present evidence that this arousal-based facilitation of self-control generalizes across emotional valences, and across self-control task domains. Finally, the pupil dilation index we describe in this work bears important advantages that may enhance the toolkit of research in the affective sciences, cognitive control and self-control.

## Materials and Methods

### Participants

Data were acquired from 43 healthy adults. Data of thirty-four participants (20 female; mean age = 22.59 ± 2.23 SD years) were included in the pupil analyses (all exclusion decisions were made based on a priori criteria that are well established in our laboratory and before analysing any data; please see the Supplementary Methods for detail). All participants were German native speakers and the experiment was conducted in German to ensure that participants were well able to follow the instructions and paradigm. Screening assured that participants were not depressed (Beck Depression Inventory (BDI) I (Beck, Steer, & Brown, 1978), German validated version by Hautzinger, Bailer, Worall, and Keller (1995)) or emotion blind (Toronto Alexithymia Scale, TAS (Bagby, Parker, & Taylor, 1994), German validated version by Franz et al. (2008)), because both conditions have been associated with altered emotion perception. Participants included in the pupil analyses on average scored 4.1 ± 2.8 SD on the BDI (cutoff for mild depression: 10). On the TAS, the mean score was 38.1 ± 8 SD (potential alexithymia: 52, cutoff: 61). With respect to the dietary health challenge task, screening assured that all participants were interested in maintaining a healthy diet, but also reported to like and consume snack foods and sweets at least on two occasions per week so that meaningful self-control challenges could be created. All participants provided written informed consent at the day of the experiment according to the Declaration of Helsinki, and the study was conducted in accordance with the regulations of the Ethics Committee of the Canton of Zurich.

### Procedure emotion task

Emotion stimuli were selected from the International Affective Picture Set (Lang, Bradley, & Cuthbert, 1999). The experimenter used the instruction detailed in Lang et al. (1999) to introduce the 9-point Self-Assessment Manikin (SAM) Scale (Bradley & Lang, 1994). Briefly, participants rated their current emotion after each trial with a version of the SAM scale validated by Suk (2006) that displayed 9 Manikins for the valence. The most negative feeling was coded as 1, and the most positive feeling was coded as 9. The scoring direction was counterbalanced across participants by randomizing whether the most negative valence would be scored on the left or on the right side of the scale. The reappraisal task was based on Wager et al. (2008). We additionally included stimuli with positive valence, which allowed us to investigate domain-general regulation and facilitates comparisons of the emotion reappraisal paradigm with the dietary health challenge task that the same cohort of participants solved on the same day (see below). Participants practiced the reappraisal task in a standardized procedure (see Supplementary Methods) on stimuli that were not repeated in the evaluated task. In the view condition, participants were instructed to simply look at the image and let the emotional response occur as elicited by the picture. Participants were asked not to modulate their emotional response, providing us with pupil data of unaltered positive and negative picture viewing (Figure 2a, dotted lines). In the reappraisal condition, participants were asked to try and come up with an alternative scenario accounting for the observed scene, such that the evoked emotional response moved more towards a neutral state.

Blocks of 20 trials of either condition were cued by displaying the words “view” or “reappraise” for 1 second centrally on the screen (Figure 1b). At the beginning of each trial, participants saw a phase-scrambled version of the stimulus image for 1 second that allowed the pupil to adapt to low-level stimulus properties. The emotion content version of the picture was then displayed for 7 seconds as in Wager et al. (2008).

Because of the adaptation period in our design, we were able to baseline-correct the signal on each trial based on the physical features of the actual stimulus displayed subsequently. This allowed us to interpret changes in pupil dilation during the view period as being primarily driven by the emotional content of the stimulus, while during reappraisal trials regulation-related pupil responses should be observed in addition to the emotional arousal pertaining to the stimuli. During the 7 seconds stimulus presentation, participants had to either view the image without regulating their emotional arousal, or reappraise their emotional responses to make them more neutral according to the trained procedure. To remind them of the currently relevant task condition, a short cue (“V” for view or “R” for “reappraise”) replaced the fixation cross overlaid on the stimulus. After the stimulus presentation, participants rated their current emotional response on the 9-point SAM valence scale (within 4 seconds). A jittered inter-trial interval (uniformly sampled from 1 to 5 seconds) separated the trials.

Block types (Reappraise Positive, Reappraise Negative, View Positive, View Negative, View Neutral) were presented in 5 pseudo-randomized orders that ensured that valence changed after each block. Each block was followed by a 15-second break, and the task was performed in a single session.

### Emotion regulation success measurement

To quantify reappraisal success, participants rated all 40 stimuli that had been presented in the reappraisal condition while sitting at a standard computer terminal.

Participants were asked to rate the images as in the “view” condition, i.e. rating the feeling elicited by the image without altering the emotion. We then quantified emotion regulation success: If negative images were successfully reappraised, participants should rate the image more positive than in the view condition. Therefore, negative reappraisal success was calculated according to Equation 1:

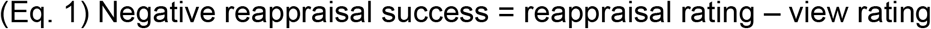

Vice versa for positive stimuli, the rating after successful reappraisal should be more negative than the view rating, and was thus calculated as in Equation 2:

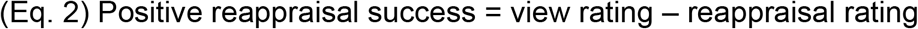

In order to obtain the overall reappraisal success score of each participant, we then averaged over positive and negative reappraisal success scores.

### Dietary health challenge task

Methodological details for this task are described in a companion paper by Maier and Hare (2020). Briefly, in the dietary health challenge task (Figure 1c), one food item was presented on each trial, and participants had 3 seconds to indicate whether they wanted to eat this food or nothing at the end of the study (participants had fasted 3 hours prior to completing this task). Choices were customized based on the individual taste and health ratings such that in challenging choices we presented each participant with foods that they had rated a) subjectively tastier and less healthy than neutral, or b) healthier and less tasty than neutral. In the remaining choices, health and taste attributes were aligned. Trial types (challenge or no challenge) were randomly mixed. The dietary health challenge success level was defined as the proportion of challenging choices during which participants refused to eat tasty-unhealthy foods or accepted to eat healthy-untasty foods.

Participants completed the dietary health challenge task and emotion regulation task in single runs with 100 trials each. Tasks were acquired in counterbalanced order, with a 7-minute break in between.

### Stimulus selection

The stimuli that participants viewed or reappraised in the emotion regulation task were selected from the International Affective Picture System (IAPS) database (for a full list please see the Supplementary Methods). These photographs have been widely used in emotion regulation research (Moser et al., 2017; Ochsner et al., 2004; Silvers et al., 2012; Wager et al., 2008) and their arousal and valence levels have been quantified and validated in large population studies (Lang et al., 1999; Lang, Greenwald, Bradley, & Hamm, 1993). For the current study, we selected IAPS Stimuli based on a validation study in a German-speaking sample (Grühn & Scheibe, 2008). Based on the mean ratings given by the young adults in this dataset, we identified 40 images that scored highest on positive and 40 images that scored highest on negative valence. To minimize confounds in identifying physiological correlates of the regulation process, we next aimed to equate the average arousal levels of these stimuli. For both positive and negative stimuli, we created two sets of 20 stimuli each. To control for the average arousal and valence levels, we distributed the images between the sets such that both sets in each domain scored on average the same for valence (mean negative = 2.25 ± 0.29 SD; mean positive = 7.25 ± 0.24 SD) and for arousal (mean negative: 6.99 ± 0.44 SD; mean positive: 2.86 ± 0.43 SD, based on the ratings of the sample in Grühn and Scheibe (2008)). We then randomly allocated for each participant which set would be presented in the “view” and “reappraise” condition for each valence domain. We also identified 20 images that scored neutral on both valence and arousal. When selecting the images, we excluded any that showed foods (due to the dietary health challenge task performed on the same day) and replaced them with the next best-scoring image.

### Pupil data acquisition

Pupil diameter was sampled at 500 Hz using an MR-compatible EyeLink II CL v4.51 eyetracker system (SR Research Ltd). Prior to the start of the paradigm, the recording quality was tested and adjusted with a spatial 9-point calibration task, covering the full screen dimensions and assuring that tracking would not be compromised by large eye-movements. During preprocessing, blinks were identified using built-in detection techniques provided by EyeLink and linearly interpolated (de Gee, Knapen, & Donner, 2014; Grueschow, Polania, Hare, & Ruff, 2015; Murphy, Vandekerckhove, & Nieuwenhuis, 2014).

Perceived brightness and stimulus contrast are low-level visual stimulus features that have previously been shown to affect pupil size (Binda, Pereverzeva, & Murray, 2013; Laeng & Endestad, 2012; Mathot & Van der Stigchel, 2015; Naber, Alvarez, & Nakayama, 2013; Naber, Frassle, & Einhauser, 2011; Porter, Troscianko, & Gilchrist, 2007; Vo et al., 2008; Wang, Boehnke, Itti, & Munoz, 2014). To facilitate isolating pupil signals related to self-control mechanisms during reappraisal, we introduced an adaption phase before each trial that allowed us to account for low-level stimulus features. One second before the actual stimulus presentation, we displayed a phase-scrambled adaptation version of the stimulus (Figure 1b, second screen) that did not reveal the semantic content of the scene, but allowed the pupil to adapt to contrast and brightness of the target stimulus (Figure 1b, third screen; please see the Supplementary Methods for details of the algorithm that matched properties of the adaptation and target stimulus pairs and Supplementary Figure 1 for the close correspondence between adaptation and target stimuli).

### Pupil analyses

Our approach applies principles of time series analysis to the pupil recordings. In order to attain a common frame of reference for the pupil dilation, all pupil time courses were z-scored per participant and run. We also baseline-corrected the within-trial pupil signal by subtracting the average pupil size during the 500 ms period before stimulus onset (Murphy, Vandekerckhove, et al., 2014). We then constructed a subtraction contrast to isolate signal components specific to regulation that differ from pupil dilation signals associated with merely viewing emotional stimuli.

We aimed at characterizing common regulation processes during both positive and negative reappraisal and thus collapsed the data over the positive and negative valence domains. To dissociate the portions of the pupil dilation signal specific to regulation, we subtracted the average pupil dilation during simple viewing of stimuli from the pupil dilation measured during reappraising equivalent content in stimuli that were on average equated for valence and arousal as described above. For convenience, we will henceforth describe this subtraction as the “Reappraise > View” contrast. By construction of these intra-individual contrasts and thereby at the same time controlling brightness between the reappraise and view conditions, we rule out confounds following the recommendations by van der Wel and van Steenbergen (2018). In line with the recommendations by Goldinger and Papesh (2012), we also baseline-corrected the signal for the tonic pupil diameter before the trial and used inter-trial intervals of at least 1 second in addition to the adaptation period. Furthermore, the Reappraise > View subtraction contrast was evaluated at the group level, and across the group, randomization ensured that both subsets of stimuli were featured equally often in the reappraise and view conditions. This entails that any stimulus-specific effects in the pupil dilation signal average out in the analysis.

To identify time periods during which the pupil dilation significantly differs between regulation and mere viewing, we performed a cluster-based permutation test following Nichols and Holmes (2002) with a cluster-forming threshold of T = 3 applied to each time bin. Briefly, we first for each participant took the mean of the reappraise time series and the mean of the view time series (collapsing across positive and negative valences) during the 7-second reappraise/view period and calculated the difference between both in order to compute the Reappraise > View contrast. We then calculated the one-sample t-statistic for each millisecond time bin in this contrast. This t-statistic indicated for each time bin whether the signal difference during reappraising versus viewing significantly differed from zero, which allowed identifying periods of consecutive time bins that exceeded the cluster-forming threshold. These defined the sizes of the temporal clusters (i.e., number of adjacent time bins exceeding the threshold of T = 3) to test against a null distribution in the next step. The null distribution was generated by permuting the labels for each time bin within-participant for 1000 iterations (flipping the sign of each time bin randomly 1000 times). On each iteration, we then again calculated for the average time series within each participant the one-sample t-statistic for each time point and identified the time bins in the permuted t-statistic vector that exceeded the cluster-forming threshold. We thereby identified the cluster sizes of each permuted cluster and stored the largest permuted cluster in the null distribution on each iteration. Note that by storing and comparing only the largest permuted cluster (and not the mean cluster size in the null distribution), we chose a more conservative approach than Nichols and Holmes (2002). To draw inference and to calculate a p-value for the temporal clusters identified in the data, we then asked how many of the randomly generated clusters from the null distribution were larger than the cluster we observed in the data. In other words, we counted how many times a cluster of the size we observed in the true data would have occurred by chance. To calculate the equivalent of a p-value, we divided this number by 1000 (iterations) in order to judge how likely it was to obtain a cluster of this size just by chance. In our case, in the real data 2190 adjacent time bins had a t-statistic greater than the cluster-forming threshold, and there were 0 clusters in the null distribution that had at least 2190 adjacent time bins with the t-statistic being greater than the cluster-forming threshold. Hence the p-value would be 0/1000, so we report it as p < 0.001. We further report the maximum t-statistic occurring in this time window and the mean value of the pupil dilation index and its standard error across the group to characterize the effect size.

Pupil dilation is governed by the sympathetic branch of the autonomic nervous system (activation / arousal) and pupil constriction is governed by the parasympathetic branch (relaxation). Therefore we assume that during the process of regulating the pupil diameter should increase and once the regulation has been successful the pupil diameter should decrease again. Arousal processes related to regulation for each individual can therefore be captured by the mean difference in the Reappraise minus View contrast during the time in which we identified significant increases in pupil dilation across all participants (between 3.4 and 5.6 seconds). In the following analyses, we will refer to this measure as Pupil Dilation Index (PDI).

### Influences on reappraisal success

We modelled emotion regulation success as a function of the pupil dilation index and the affective distance to be regulated, using a Bayesian linear regression model described in Equation 3a below:

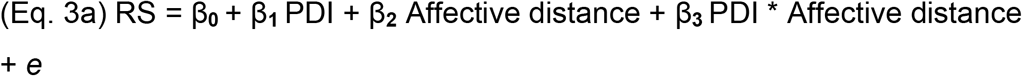

Where *PDI (pupil dilation index)* denotes for each participant the mean difference in the pupil dilation signal during the significant regulation time window in the contrast Reappraise > View. *Affective distance* describes the average regulation distance to be covered for each participant, measured as the absolute distance of the view ratings for the regulated images from neutral. We included an interaction term for PDI and affective distance because it may be easier or harder to regulate depending on the regulation starting point. *Reappraisal Success* was measured as the average of positive and negative reappraisal success for each participant that were separately scored according to Eq. 1 and 2.

In order to control for potential confounds that may alternatively explain regulation success, we augmented the model as described in equation 3b:

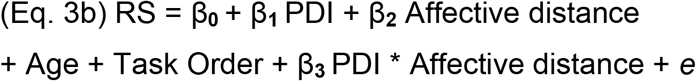

Here, in addition to the coefficients described in equation 3a, the model included a standardized and mean-centred term for the age of each participant, and a factor that indicated in which order the emotion and dietary health challenge tasks were completed.

### Out-Of-Sample Predictions

To test the ability of the main model described in Eq. 3a to predict out of sample, we used a leave-2-samples-out approach. We generated all 561 possible combinations of training/test sets: in each training set, we estimated the model on the data of 34 participants and predicted for the test set (the two left-out participants), which of these two individuals would be more successful at regulating. We then compared the predicted to the true regulation success ranking for each of these 561 combinations and calculated the accuracy of our predictions, i.e. the proportion of true predictions out of all 561 predictions made. In order to quantify how often this accuracy would occur by chance, we generated a null distribution of prediction accuracies from 1000 iterations running the model on random training data. To this end, on each iteration, we permuted the labels for the regulation success scores of the training sets (randomly multiplying half of the training set success scores by -1) and trained the model on these random outcome values. Based on these estimates, we then again predicted for all possible combinations of training/test sets which of two left-out participants regulated more successfully. On each iteration, we calculated the accuracy of the prediction (i.e. how many pairs of participants were correctly predicted out of all possible combinations). We thereby obtained a distribution of 1000 accuracies that would have occurred by chance. To calculate a p-value, we computed how often accuracies greater than the one that was obtained from predictions based on the true data would occur in this null distribution (i.e., by chance), and then divided this value by 1000 to account for the number of iterations.

### Predictive validity across self-control domains

In order to test the predictive validity of the pupil dilation index across self-control domains, we calculated a Bayesian rank correlation between the pupil dilation index from the emotion regulation task and the health challenge success level achieved by the same individuals in a separate dietary health challenge task measured on the same day. To assure that any effects were independent of the order in which tasks are performed, we modelled health challenge success controlling for task order using the Bayesian linear regression model described in Eq. 4:

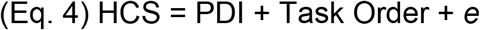

Where *HCS* denotes the overall health challenge success in the dietary self control task, operationalized as the percentage of challenge trials during which participants refused unhealthy-palatable foods or accepted to eat healthy-unpalatable foods, *PDI* is the pupil dilation index, and *Task Order* is a factor accounting for the order in which the emotion regulation and dietary health challenge tasks were performed.

### Statistical Analyses

All analyses were performed with the R (R Core Team, 2013) and Matlab (The MathWorks Inc., 2012) statistical software packages. For all Bayesian modelling analyses, we used the default, uninformative priors specified by the brms (Bürkner, 2017) or BEST (Kruschke, 2013) R-packages. The use of uninformative priors entails that our Bayesian analyses yield results very similar to those of conventional frequentist statistics. Results from all Bayesian analyses are reported as the mean of the posterior predictive distribution that indicates the most credible estimate, along with its standard deviation (SD) and the 95% Highest Density Interval that denotes the range in which 95% of the credible estimates fall. This is denoted as 95 % Credible Interval (CI). The notation PP() indicates the posterior probability that the relation stated within the parentheses is true.

## Code and data availability

Code and data for the presented analyses will be made openly available upon publication of the paper on OSF. During review, it will be made available to reviewers upon request.

## Acknowledgements

This work preprinted on bioRxiv is a companion paper to work by Silvia Maier and Todd Hare that tested a separate hypothesis in the same dataset. The prior work did not make use of the pupillometry data and those data have not been reported before. For the present work, S.U.M. and M.G. contributed equally to designing the study, analysing the data and writing the manuscript;

S.U.M. conducted the experiments. The authors thank Astrid Dobler and Jonathan Schaffner for assistance with data collection and documentation. We gratefully acknowledge funding through EU FP7-KBBE Grant 607310 and SNF Grant CRSII5_177277 (S.U.M.), and the Richard Büchner Foundation (M.G.). The funders had no role in the conceptualization, design, data collection, analysis, decision to publish, or preparation of the manuscript.

## Competing interests

S.U.M. and M.G. both report no financial or non-financial competing interests.

## Supplementary Information

### Supplementary Figures

**Supplementary Figure 1.**
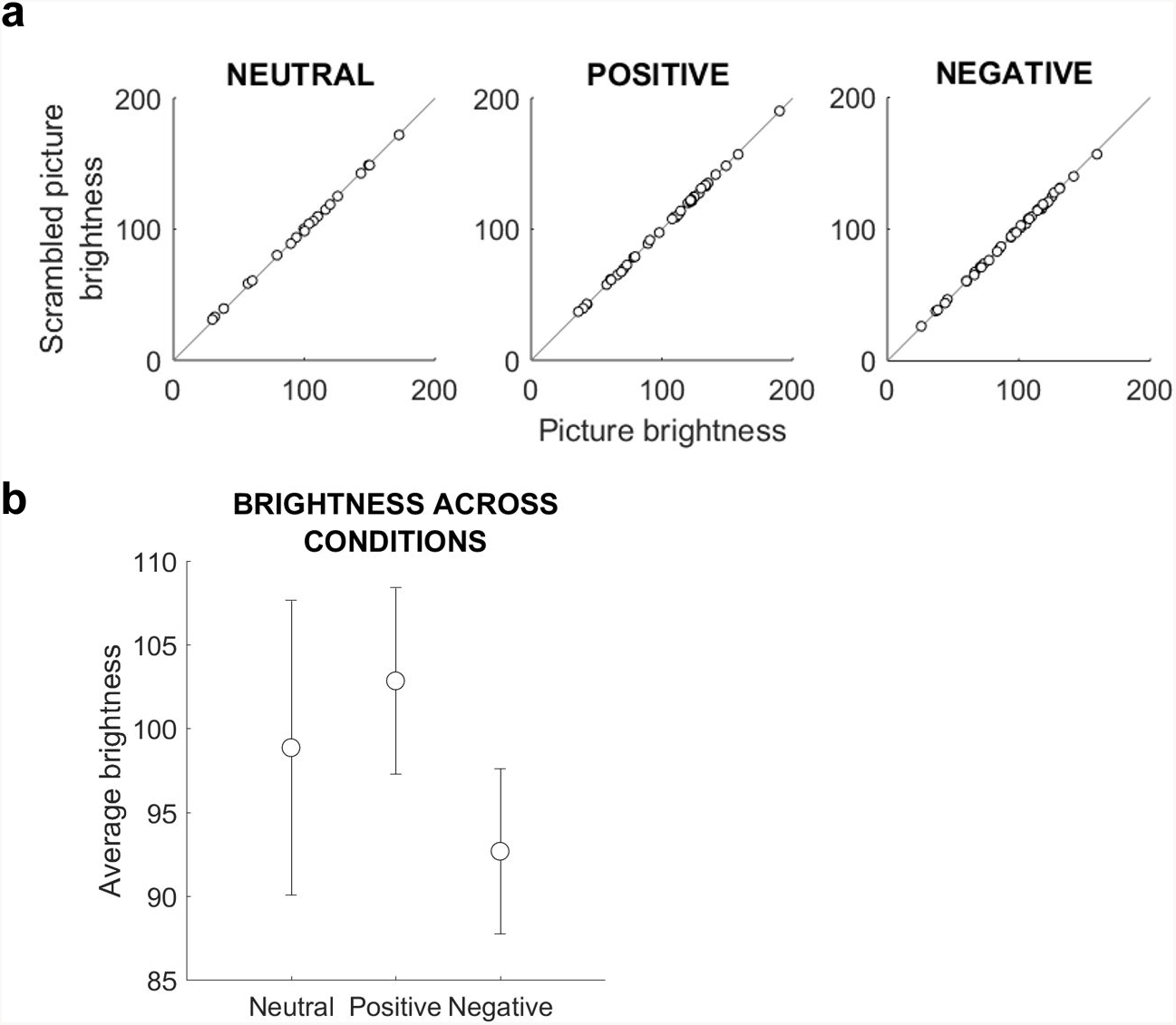
Brightness matching of adaptation and target stimuli. Panel **a)** depicts the brightness-match of the adaptation and target stimuli for the neutral, positive and negative blocks. Each dot represents one stimulus. The 45-degree line is added for visual evaluation, indicating identity between the brightness of the scrambled adaptation stimuli and the target stimuli in which the content was visible. Panel **b)** plots the mean brightness (dots) and the standard error of the mean (bars) for the stimuli in the neutral, positive and negative stimulus sets.

**Supplementary Figure 2.**
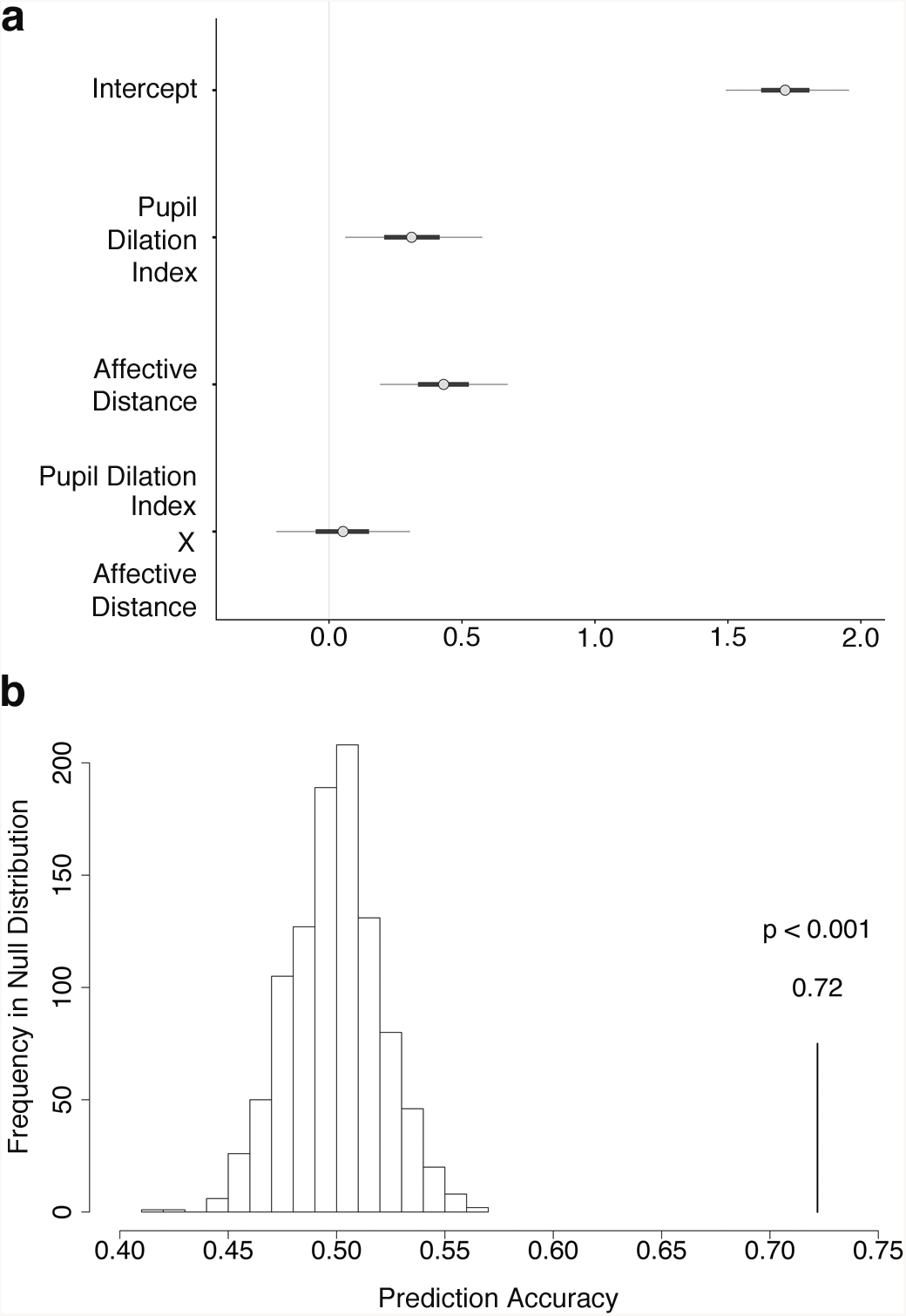
Results from the control analysis for regulation success prediction by blocks. Panel **a)** depicts the coefficients of the Bayesian linear regression fit of the model given in Eq. 3a for the control analysis using measures of reappraisal success and affective distance that were constructed from the view ratings given for the equivalent set of stimuli in the view block that had never been reappraised. As in the original model, we observed that regulation success increased both with greater Pupil Dilation Index and Affective Distance. The plot shows the mean beta estimates (grey dots) as well as the range of coefficients within the 90% Credible Interval that is represented by the light grey horizontal line (thick horizontal bars = 50% Credible Interval). Panel **b)** shows the corresponding out-of-sample prediction. The results from above were cross-validated with a Leave-2-Participants-Out approach. Based on data from N-2 participants to which the above model was fit, we predicted which of the two left-out individuals regulated more successfully. The model predicted with 72% accuracy significantly above chance (p < 0.001).

### Supplementary Methods

#### Dataset

The pupil data we report here were recorded while acquiring an fMRI dataset that has first been reported in a companion paper by Maier and Hare (2020). There, the authors tested an unrelated hypothesis using the fMRI and behavioural portions of the dataset. The present manuscript is the first report of the pupillometry data.

#### Participant exclusion

Out of the 43 collected datasets, the following datasets could not be evaluated for the pupil analyses: 7 participants did not complete the emotion reappraisal paradigm with sufficient data quality (five fell asleep during longer stretches of the task (detected by the eye-tracker), one deliberately closed the eyes when negative stimuli were displayed, and one reported experiencing pain due to head positioning during the task. We reasoned that this participant who reported his discomfort only after the study likely engaged in constant self-control that would interfere with our analyses). These participants were also excluded from the analyses in the companion paper. Pupil data could not be evaluated for two additional datasets because the eye-tracker did not record the start of the experiment correctly.

#### Stimulus sets

Negative Set A: 1300, 2055.1, 2095, 2981, 3015, 3181, 3301, 3550, 6020, 6212, 6370, 6540, 6838, 9040, 9180, 9181, 9265, 9435, 9520, 9570; Negative Set B: 1525, 2352.2, 2683, 2800, 3051, 3250, 3530, 6312, 6415, 6570.1, 9140, 9250, 9252, 9253, 9300, 9430, 9561, 9571, 9635.1, 9800; Positive Set A: 1460, 1710, 1721, 1750, 1810, 1920, 2050, 2080, 2091, 2224, 2260, 2311, 2351, 2375.2, 2550, 5600, 5626, 5831, 5890, 8190; Positive Set B: 1440, 1463, 1720, 1731, 2058, 2071, 2150, 2170, 2303, 2345, 2620, 2655, 2660, 5390, 5594, 5628, 5830, 7580, 8461, 8497; Neutral Set: 2020, 2200, 2214, 2357, 2480, 2493, 2570, 2880, 2890, 7030, 7036, 7150, 7161, 7170, 7186, 7224, 7590, 7705, 7830, 8030.

#### Reappraisal task instructions

To practice the reappraisal method, we provided standard written instructions that contained one example for positive and negative pictures before participants practiced modulating their emotional responses. Negative pictures for example contained scenes of humans suffering from wounds, war or crime scenes, or dead bodies of humans or animals. Positive scenes displayed animal or human babies, individuals playing or laughing, or nice views of landscapes. Participants were asked to modulate their interpretation of the pictures content such that negative feelings should become less negative, and positive feelings less positive. For example, one could think of the image as a scene or mock-up from a movie: Things are not as bad or good as they seem, but just staged. After receiving these standard instructions, participants trained with a computerized version of the task to familiarize themselves with the display sequence for one trial (adaptation stimulus – revealed stimulus – SAM rating screen), as it would be depicted in the actual experiment. First, participants reappraised one positive and one negative picture (order counterbalanced) at their own pace, and then two more images while picture presentation and emotion rating were presented with free timing and timed as in the actual experiment. Training was performed on pictures that were not presented in the actual experiment later. We assured participants felt competent to use the procedures before we started the recording as judged by mutual agreement between the participant and the experimenter.

#### Basic sanity check for emotion reappraisal success

Behavioural and fMRI data of this experiment were first reported in Maier and Hare (2020). For completeness and accessibility, we reproduce in this section the analysis approach for reappraisal success that has first been reported in the companion paper.

The linear regression model summarized by Equation S1 (given in condensed notation) was estimated to test whether ratings differed significantly between the view and reappraise conditions:

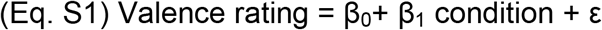

On the trial-level, the variable *valence rating* denoted the rating given on the respective trial. It was coded from 1 (very sad) to 9 (very happy) in steps of 1.

*Condition* was modelled as a factor (levels: 0 = neutral view, 1 = negative view, 2 = negative reappraisal, 3 = positive reappraisal, 4 = positive view). The model included subject-specific random intercepts and slopes for the condition.

#### Control analysis for the reappraisal success measure

The “view” ratings of the reappraised images were collected after the scan session, i.e. after reappraising the content in the scanner. Previous work in the literature suggests there may be spillover effects from reappraisal in neural signals when the same stimuli are presented again (MacNamara et al., 2010), although these effects may be relatively transient and stronger when stimuli are reappraised multiple times (Denny, Inhoff, Zerubavel, Davachi, & Ochsner, 2015). As a sanity check to exclude that any potential spillover effects from regulation would change our conclusions, we ran a control analysis following our method laid out before, but relying on the view ratings for the equivalent set of stimuli that were not reappraised but only viewed during the task.

Analogous to Equation 1, for the negative stimuli, we calculated for each participant according to Equation S2:

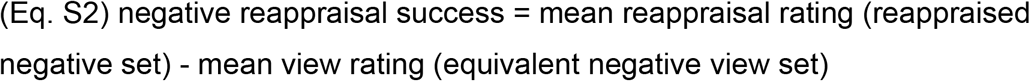

Analogous to Equation 2, for the positive stimuli, we calculated for each participant according to Equation S3:

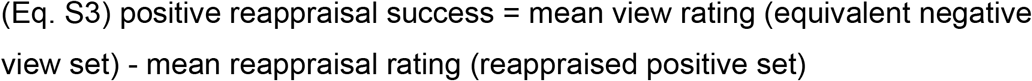

To obtain the overall *regulation success score*, we then again took both success scores and calculated the mean across both positive and negative valence.

*Affective Distance* was also constructed using the view ratings from the view stimulus set for this control analysis. For each of the equivalent positive and negative stimulus sets we first averaged the mean difference of the view ratings from neutral, and then calculated the mean over the negative and positive sets.

With these measures, we then repeated the analysis described in Eq. 3a as well as the out-of-sample crossvalidation based on these regulation success measures.

### Stimulus construction for the pupil adaptation phase

Low-level feature matching was achieved via two image-processing steps. For any given trial, we first phase-scrambled the spatial frequency information of the target image (180°) using identical procedures as previously reported in Rieger et al. (2013). This approach renders the semantic content of the picture unrecognizable, while retaining the global contrast of the picture. The 180° phase-scrambled version of the target picture served as our adaptation stimulus. Such phase-scrambling methodology has previously been successfully employed to dissociate sensitivity to contrast and spatial frequency in human primary visual cortex (Rieger et al., 2013). This step matched the contrast between the adaptation and target stimulus. This technique typically yields pictures that may appear darker than the original. Therefore the target image was brightness-adjusted (typically decreased by 10%) relative to the phase-scrambled version of the image using Matlab functions provided by Chris Rorden (https://www.mccauslandcenter.sc.edu/crnl/tools/bmp_contrast) and the Matlab Image Processing Toolbox. The result of the second image-processing step is thus equal global brightness distribution between the adaptation image and the target image, which contains semantic content to either be viewed or reappraised. Stimuli were presented at 126.5 cm viewing distance on a mean grey projection screen, with a stimulus size of 10.9 width x 8.1 cm height. Our procedure successfully matched the contrast and brightness for each pair of adaptation and target stimuli (Supplementary Figure 1), where plotting the brightness properties of the adaptation against the target stimuli results in almost perfect identity in brightness features.

#### Control of visual stimulus properties

Across positive, negative and neutral emotion conditions, our stimulus sets may still differ in their low-level stimulus properties, as we applied the matching algorithm only within-trial. Indeed, we observed that on average, the pupil restricted more during the positive conditions than the negative (Figure 2A). In line with Henderson et al. (2014), this may be an effect of anticipating a threat in the negative condition, as stimuli were presented in blocks. The pupil also stayed on average more constricted throughout the positive view and regulation trials, which may be due to brightness differences of the stimuli. Although there was no difference at the significance level of p < 0.05, the average brightness for the positive stimuli tended to be higher than the average brightness of negative stimuli (Supplemental Fig. S1b; mean brightness positive = 102.85 cd/m^2^, mean brightness negative = 92.67 cd/m^2^, p = 0.18, 95% CI = [-4.80, 25.15]).

However, we control for any remaining differences with our design. In our main analysis of interest, we calculate a contrast between regulation and viewing that averages regulation and view signals over both positive and negative valence domains. Therefore, any brightness-induced differences in pupil size cancel out in this contrast, as both positive and negative stimuli appear on both sides of the subtraction, in the regulate or view condition.

Hence the remaining differences are related to regulation, not the physical stimulus properties.

#### Statistical packages

All Bayesian analyses were performed with the R (R Core Team, 2013), STAN (Carpenter et al., 2016) and JAGS (Plummer, 2003) statistical software packages. Bayesian regressions were run using the brms package (Bürkner, 2017) that is an interface between R and STAN. All correlations (Kruschke, 2015) and t-tests were computed using Bayesian Markov Chain Monte Carlo (MCMC) sampling methods using R in combination with JAGS (Kruschke, 2013). The behavioural plots in Figure 1 were created using the yarrr package (Phillips, 2017), the scatter plot in Figure 3C was created with ggplot (Wickham, 2017). The package pracma (Borchers, 2018) was used for data handling.

### Supplementary Results

#### Emotion ratings with and without reappraisal

For the reader’s convenience, we re-iterate as a sanity check in this paragraph the behavioural results that are reported in a companion paper by Maier and Hare (2020). In brief, the reappraisal task worked as expected. On the scale from 1 (very negative) to 9 (very positive), ratings after reappraising negative content shifted closer to neutral (mean negative reappraise rating = 4.25 ± 0.81 SD) and were distinctly more positive than ratings after viewing negative content as assessed by the Bayesian equivalent of a paired T-test (BEST (Kruschke, 2013)) that estimated the Posterior Probability of Negative Reappraise being greater than Negative View ratings (PP(Negative Reappraise > Negative View Ratings)) > 0.9999. Likewise, current emotions were rated more neutrally after reappraising the positive stimuli (mean positive reappraise rating = 5.21 ± 0.9 SD). Ratings after positive reappraisal were also clearly less positive than when participants viewed positive content and let their emotional response occur naturally (PP(Positive Regulate < Positive View Ratings) > 0.9999).

Thus, participants successfully regulated their emotions by reappraising the stimulus content.

#### Control analysis for the reappraisal success model

We quantified the emotion regulation success level mathematically based on the procedure by Wager et al. (2008). In order to exclude confounds due to potential spillover effects of reappraisal (MacNamara et al., 2010), we repeated the analysis described in Eq. 3a using measures for emotion reappraisal success and affective distance that were constructed block-wise, based on ratings given for the equivalent stimulus set used in the “view” condition. These measures were thus based on the mean view ratings from the equivalent negative and positive stimulus sets that had been equated with the reappraised stimulus sets for average arousal and valence. Like the reappraised images, these images in the view block had been viewed in the scanner for the first time.

Using these measures, we found the same pattern of results as before (Supplementary Figure 2; Supplementary Table 2): the pupil dilation index explained a substantial portion of the reappraisal success (beta = 0.31 ± 0.16 SD, 95% Credible Interval (CI) = [0.01; 0.63]), above and beyond the effects of affective distance (beta = 0.43 ± 0.15, 95% CI = [0.13; 0.73]). The crossvalidation for this control analysis also corroborated our previous results (Supplementary Figure 2; prediction accuracy = 72%, p < 0.001). Hence our conclusions based on the more fine-grained, stimulus-wise constructed regulation success measure replicated.

### Supplementary Discussion

#### Development and rationale of the pupil dilation index

We determined the time window during which we identified regulation signals by calculating a cluster-based permutation t-test to separate the pupil signal related to regulating from the signal related to solely viewing emotional content without regulation demands. While this approach aims at incorporating all signals that relate to the regulation process, the within-participant subtraction (van der Wel & van Steenbergen, 2018) of “regulate” and “view” signals renders the remaining pupil dilation index robust and ensures that only regulation-relevant parts of the pupil dilation are evaluated in order to determine the timing.

Because our design equated the stimulus sets between view and regulate conditions for their average valence and arousal, we thereby only evaluated those portions of the dilation response that go beyond the average dilation that was observed when emotional stimuli with similar properties were contemplated. Following the reasoning of Hess and Polt (1964), who suggested that “total mental activity” could be captured as a combination of the amplitude and latency of the pupil dilation response, we then calculated the mean difference of the pupil dilation amplitude during the significant regulation time between the reappraisal and view condition to isolate the regulation-related components and determine the pupil dilation index we describe in this work (also see Kinner et al. (2017)). This index thus captures all processes that are directly relevant to regulating.

#### Potential reappraisal spillover

Previous work has shown that reappraisal may under certain conditions spill over on subsequent processing and rating of the stimuli. MacNamara et al. (2010) used standardized reappraisals that negatively framed unpleasant and neutral stimuli and found that these affected neural signals and ratings when presenting the same stimuli again 30 minutes later. However, the effects were more pronounced for neutral than negative stimuli. A study by Denny et al. (2015) presented negative stimuli in different scan sessions one day after first reappraisal and again a week later in a view or reappraise condition. In the reappraise condition, participants were to make their negative feelings less negative. Their results suggested that amygdala responses to negative stimuli were only attenuated in the long term after repeated reappraisal, but not when negative stimuli were only reappraised once. A study by Walter et al. (2009) found effects of reappraisal compared to viewing aversive stimuli that lasted up to 10 minutes after emotion regulation, and also suggested changed memory encoding when tested a year later (Erk, Von Kalckreuth, & Walter, 2010). In summary, reappraisal effects might be relatively transient and stronger when stimuli are reappraised multiple times. In our experiment, participants re-rated the stimuli outside the scanner ca. 30-45 minutes after completing the last reappraisal session. To rule out potential concerns, we checked our results in a control analysis with measures constructed based on the “view condition” (i.e., an equivalent set of stimuli that had not been reappraised), and found that both the regression model as well as the out-of-sample cross-validation corroborated our previous results, ruling out reappraisal spillover as a main determinant of our results.

### Supplementary Tables

**Supplementary Table 1.**
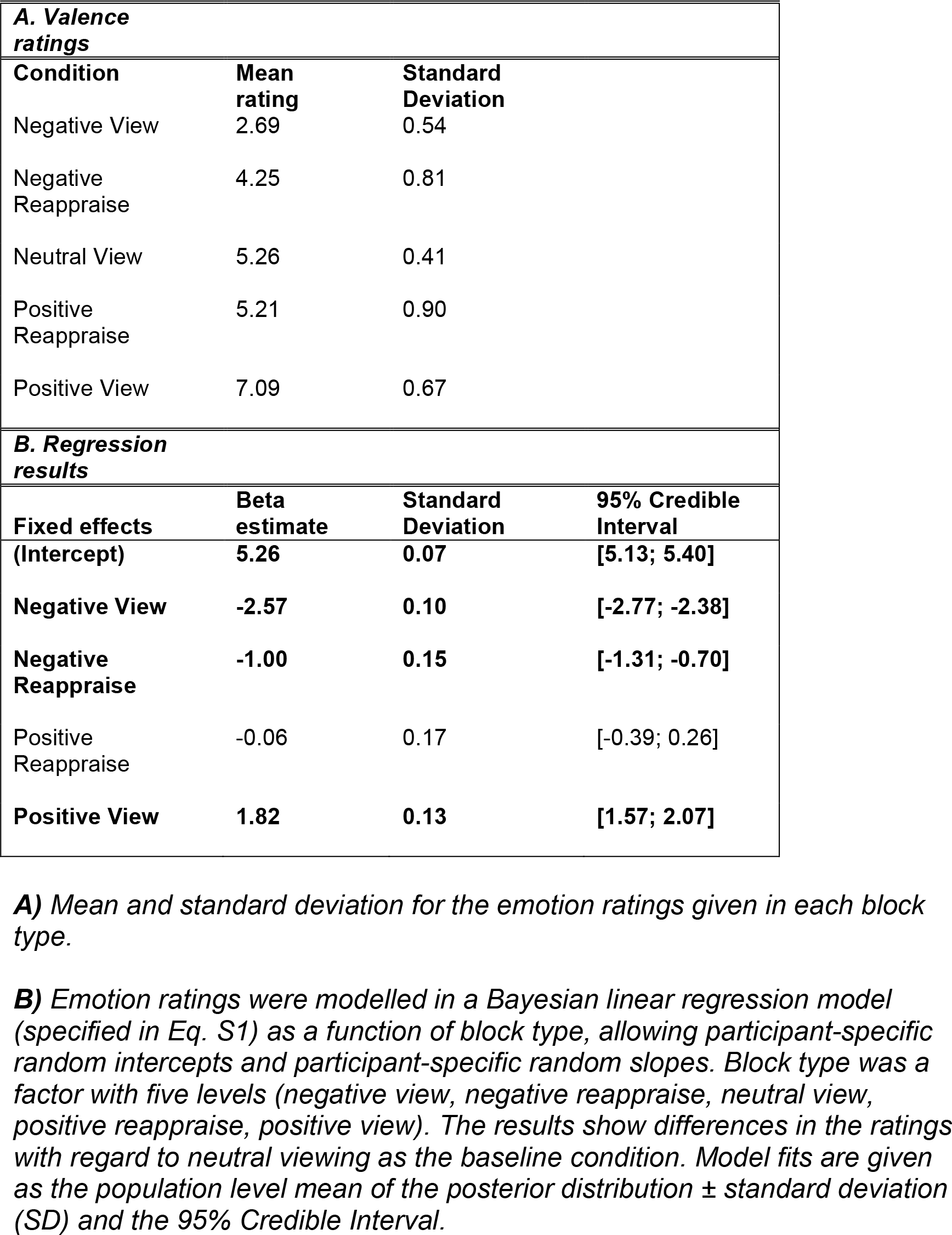
Emotion ratings by condition. (Results in this table are reproduced from Maier and Hare (2020))

| **A. Valence ratings** |  |  |  |
| --- | --- | --- | --- |
| **Condition** | **Mean rating** | **Standard Deviation** |  |
| Negative View | 2.69 | 0.54 |  |
| Negative Reappraise | 4.25 | 0.81 |  |
| Neutral View | 5.26 | 0.41 |  |
| Positive Reappraise | 5.21 | 0.90 |  |
| Positive View | 7.09 | 0.67 |  |
| **B. Regression results** |  |  |  |
| **Fixed effects** | **Beta estimate** | **Standard Deviation** | **95% Credible Interval** |
| (Intercept) | 5.26 | 0.07 | [5.13; 5.40] |
| Negative View | -2.57 | 0.10 | [-2.77; -2.38] |
| Negative Reappraise | -1.00 | 0.15 | [-1.31; -0.70] |
| Positive Reappraise | -0.06 | 0.17 | [-0.39; 0.26] |
| Positive View | 1.82 | 0.13 | [1.57; 2.07] |

**A)** Mean and standard deviation for the emotion ratings given in each block type.
**B)** Emotion ratings were modelled in a Bayesian linear regression model (specified in Eq. S1) as a function of block type, allowing participant-specific random intercepts and participant-specific random slopes. Block type was a factor with five levels (negative view, negative reappraise, neutral view, positive reappraise, positive view). The results show differences in the ratings with regard to neutral viewing as the baseline condition. Model fits are given as the population level mean of the posterior distribution ± standard deviation (SD) and the 95% Credible Interval.

**Supplementary Table 2.** Control analysis: emotion regulation success score and affective distance calculated from block of not reappraised images.

|  | **Beta Estimate** | **Standard Deviation** | **95% Credible Interval** |
| --- | --- | --- | --- |
| **(Intercept)** | **1.72** | **0.14** | **[1.44; 2.00]** |
| **Pupil Dilation Index** | **0.31** | **0.16** | **[0.01; 0.63]** |
| **Affective Distance** | **0.43** | **0.15** | **[0.13; 0.73]** |
| Pupil Dilation Index X Affective Distance | 0.05 | 0.16 | [-0.25; 0.36] |

*Results from the Bayesian Linear Regression specified in Eq. 3a. In this control analysis, we used the “view” condition blocks of stimuli that were not reappraised to construct the regulation success score and affective distance measures.*

*The emotion regulation success score was calculated as follows:*

*Analogous to Eq. 1, for the negative stimuli, we calculated for each participant their average rating after regulation in the negative condition and then subtracted their average rating after viewing the other, equivalent negative picture set that was presented in the scanner for mere viewing.*

*Analogous to Eq. 2, for the positive stimuli, we calculated for each participant the average rating for the other, equivalent positive picture set that was presented in the scanner for mere viewing and subtracted their average rating after regulation in the positive condition.*

*We then again took both success scores and calculated the mean regulation success score across both valences.*

*Affective Distance was constructed as well from the equivalent, non-reappraised view blocks. We first calculated for each of the positive and negative stimulus sets the mean difference of the view ratings from neutral, averaged it, and then computed the mean over the negative and positive sets.*

*The regulation success score was modelled by the mean-centred and standardized coefficients for Pupil Dilation Index (measured as mean difference in the pupil dilation curve for the Regulate > View contrast in the significant regulation time between 3.4 and 5.6 seconds) and Affective Distance. Pupil Dilation Index and Affective Distance were interacted, to test whether it is more or less effortful to regulate with smaller or greater affective distances. Model fits are given as the population level mean of the posterior distribution ± standard deviation (SD) and the 95% Credible Interval.*

*As this table shows, compared to Table 1, the results and thus our conclusions remain qualitatively unchanged.*

## Notes

### Competing Interest Statement

The authors have declared no competing interest.

